# Shape-related characteristics of age-related differences in subcortical structures

**DOI:** 10.1101/232439

**Authors:** Christopher R. Madan

## Abstract

**OBJECTIVES:** With an increasing aging population, it is important to gain a better understanding of biological markers of aging. Subcortical volume is known to differ with age; additionally considering shape-related characteristics may provide a better index of age-related differences in subcortical structure. Recently fractal dimensionality has been shown to be more sensitive to age-related differences, but this measure is borne out of mathematical principles, rather than quantifying a neurobiologically relevant characteristic directly. We considered four distinct measures of shape and how they relate to aging and fractal dimensionality: surface-to-volume ratio, sphericity, long-axis curvature, and surface texture.

**METHODS:** Structural MRIs from two samples, with a combined sample size of over 600 healthy adults across the adult lifespan, were used to measure age-related differences in the structure of the thalamus, putamen, caudate, and hippocampus. For each structure, volume and fractal dimensionality were calculated, as well as each of the four distinct shape measures. These measures were then examined in their utility in explaining age-related variability in brain structure.

**RESULTS:** The four shape measures were able to account for 80-90% of the variance in fractal dimensionality, indicating that these measures were sensitive to the same shape characteristics. Of the distinct shape measures, surface-to-volume ratio was the most sensitive aging biomarker.

**CONCLUSION:** Though volume is often used to characterize inter-individual differences in subcortical structures, our results demonstrate that additional measures can be useful complements to volumetry. Our results indicate that shape characteristics of subcortical structures are useful biological markers of healthy aging.

## 1. Background and Objectives

As the world’s aging population continues to increase, it is important to gain a better understanding of biological markers of aging. A variety of markers have been found to be useful in this regard—including epigenetic, physiological, neuroanatomical, and cognitive measures (Bae et al. 2013; Chen et al., 2015; Hannum et al., 2013; Horvath, 2013; Reagh & Yassa, 2017; Salthouse, 2011; Small et al., 2011; Walhovd et al., 2011). With respect to the brain, it is well established that there are age-related differences in the volume of subcortical structures (Allen et al., 2005; Goodro et al., 2012; Inano et al., 2013; Long et al., 2012; Potvin et al., 2016; Raz et al., 2005; Tamnes et al., 2013; Walhovd et al., 2005, 2011; Yang et al., 2016). However, it is important to acknowledge that volume is a summary statistic of the three-dimensional segmented structure and that it may be neglecting other facets of the structure that also vary with age, such as morphological (i.e., shape-related) characteristics. More directly, it is relatively unlikely that volumetric changes in subcortical structures would change without concurrent changes in the shape of the structure—that is, for a structure to maintain the same general form and merely ‘scale’ in size. As such, any inter-individual characteristic associated with volumetric differences, such as aging or neurodegenerative diseases, would likely be identified by simultaneously considering both volumetric and morphological properties (additional measures, such as neuropsychological tests and genetic risk factors would also be beneficial). It is an open question, however, as to what measure could be used along with volume to characterize these morphological properties, which are also neurobiologically relevant. Here we sought to examine the sensitivity of different morphological measures in indexing healthy age-related differences in subcortical structures and serving as more robust neuroanatomical markers of aging.

A recent study by Madan and Kensinger (2017a) suggested that fractal dimensionality, a measure of structural complexity, might be such a measure. In their study, fractal dimensionality indexed age-related differences better than volume, corrected for intracranial volume (i.e., ICV-corrected). Fractal dimensionality measures the volumetric properties across different spatial scales (i.e., resolution; see Figures 1 and 2 of Madan & Kensinger, 2016), allowing for a scale invariant calculation of morphological characteristics. This measure was found to be generally more sensitive to age-related variability in the subcortical structures than volume—it has been demonstrated to be a useful mathematical approach to characterizing complex structures in many domains (Di Ieva et al., 2014, 2015; Lopes & Betrouni, 2009). However, it is unlikely that fractal dimensionality is directly related to neuroanatomical changes—that is, the brain is not changing in fractal dimensionality with age, but rather that there are not-yet-understood systematic changes that fractal dimensionality is sensitive to detecting. If we accept that subcortical structures vary in volume in relation to aging, one must consider how this occurs within the brain as constrained by biology. If the thalamus is decreasing in volume due to age atrophy, it cannot simply ‘scale’ in-place while keeping the same relative shape. First, subcortical structures share boundaries with other structures—gaps do not appear throughout the brain due to these volumetric decreases— so the shape of structures must be inter-related. Second and relatedly, it is likely that the large-scale structural properties of these subcortical structures must also change in their broad curvature.

**Figure 1.**
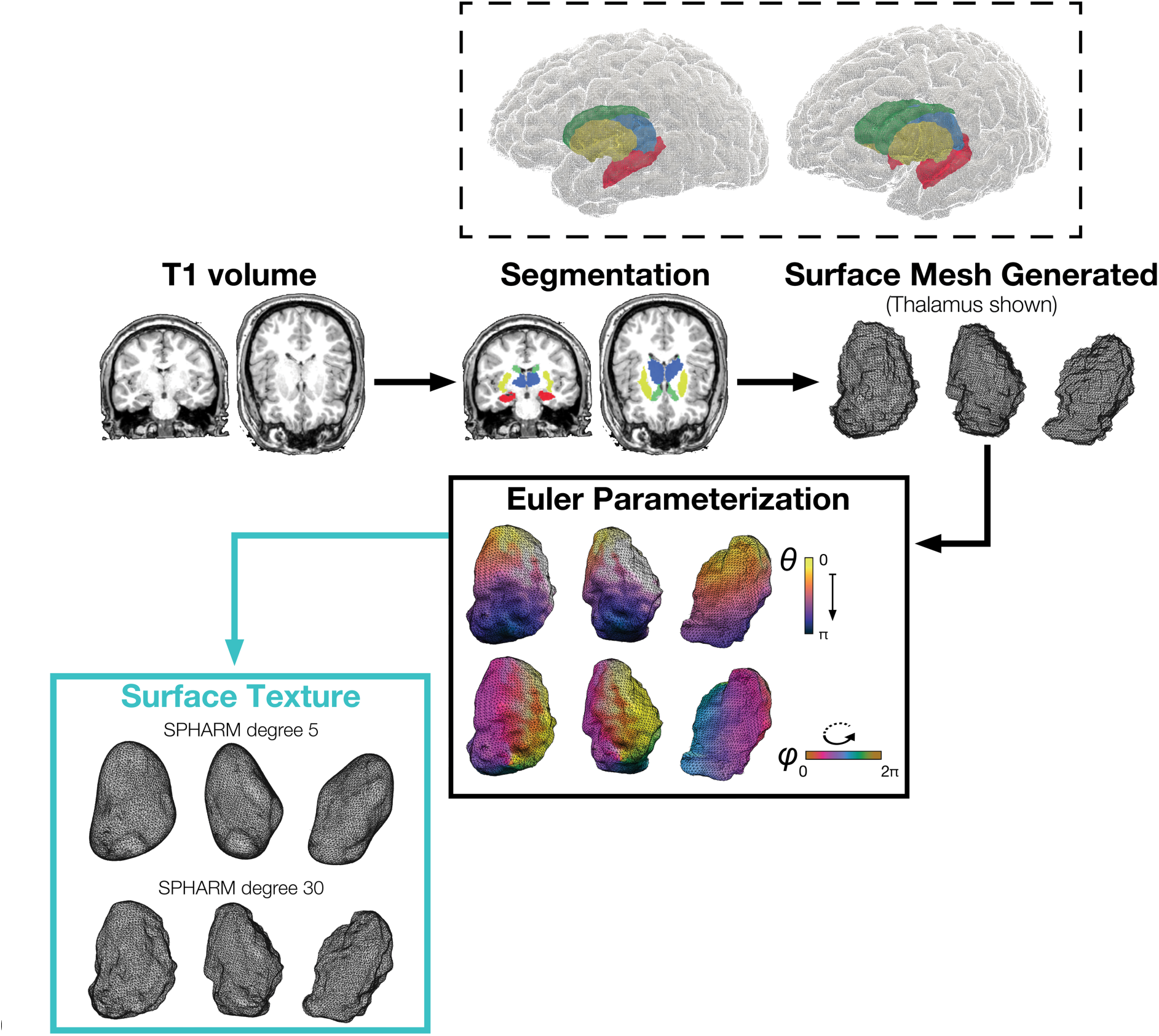
Illustration of the process used to calculate the surface mesh and texture. Glass brain 3D reconstruction constructed based on Madan (2015).

**Figure 2.**
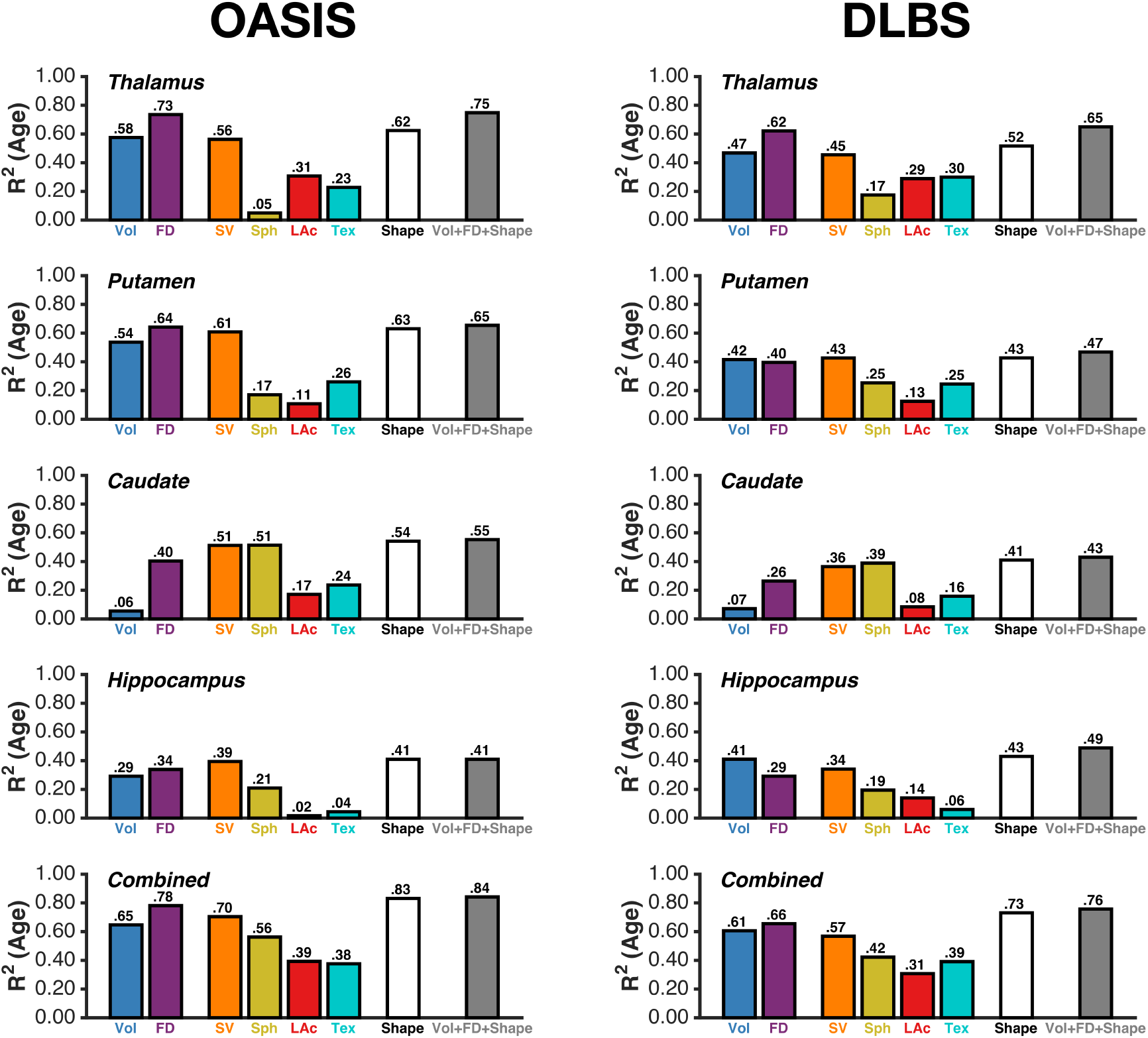
Variance explained in age, for each of the structures, morphological measures, and samples considered. ‘Shape’ is an aggregate of SV, Sph, LAc, and Tex. Key: Vol, volume; FD, fractal dimensionality; SV, surface-to-volume ratio; Sph, sphericity; LAc, long-axis curvature; Tex, shape texture; see Table 1 for additional details and comparisons.

Examining the differences in explained variability (*R^2^*) reported in Madan and Kensinger (2017a, Figure 2) for volume and fractal dimensionality, as well as the relationship between volume and fractal dimensionality (Madan and Kensinger, 2017a, Figure 5) it appears that fractal dimensionality is particularly beneficial, beyond volume, in measuring age-related differences in the structure of the thalamus, putamen, caudate, and hippocampus (see Figure 1 for visualizations of these structures). Here we consider four measures that would each be indexed by fractal dimensionality, but would not be detected by volume: surface-to-volume ratio, sphericity, long-axis curvature, and surface texture. Each of these discretizes shape-related information based on the relative scale of potential structural complexity characteristics.

1. The *ratio of surface area to volume* can be used as a coarse measure of a structure’s compactness and has long been used in characterizing the properties of 3D structures (i.e., stereology) (Lewis, 1976; Weibel et al., 1966). This ratio value will be relatively small for compact structures, but will be markedly larger for a structure that is more flattened or otherwise spread out.
2. *Sphericity,* a measure of how closely a shape resembles a sphere, measured as the ratio of the surface area of a sphere with the same volume as the structure, relative to the actual surface area of the structure (Wadell, 1932, 1935; Wentworth, 1933).
3. *Long-axis curvature* was measured by first determining the ‘mean meridian’, a curved line that went through the central mass of the structure, and has a long-standing history in the characterization of biological structures (Blum, 1973; Yushkevich et al., 2006, 2007). Long-axis curvature was operationalized as the ratio between the lengths of a curved line (spline) that travels along the mean meridian of the structure, connecting the most extended ends of the structure and traveling through the central mass of the structure, and a line that connects the two ends of the structure using the shortest straight-line distance.
4. A remaining morphological feature is the *surface texture* or roughness of the structure. This measure would correspond higher-frequency in the structure’s shape and has previously been investigated in relation to fractal dimensionality in other fields of research (e.g., Gârding, 1988; Lespessailles et al., 2006; Lopes et al., 2011; Pentland, 1985; Sarker & Chaudhuri, 1992; Thomas et al., 1999). Here we quantified the surface texture of a structure by reconstructing the subcortical structure’s topological frequency using spherical harmonics (SPHARM) (Chung et al., 2008; Gerig et al., 2001a,b; Madan & Kensinger, 2017b; Shen et al., 2007, 2009), based on Fourier series mathematics. SPHARM has also been related to the fractal dimensionality of brain structures (Madan & Kensinger, 2017b; Yotter et al., 2011). By comparing the surface area between SPHARM surfaces with differing maximum numbers of degrees we can measure the surface texture (roughness) of structures. This ratio is essentially a comparison between the surface area of a smoothed version of the structure that nonetheless captures the global shape, relative to the surface area of a mesh that does capture the nuances and local features of the structure. The difference between these two sets of coordinates effectively represents a ‘displacement map’ (Blinn, 1978; Lee et al., 2000).

By characterizing these distinct morphological measures of subcortical structures, we sought to both attain a better understanding of the shape-related features that were indexed by fractal dimensionality, as well as potentially determine a more precise measure of morphology that is further sensitive to as a neuroanatomical marker of aging. Here we evaluated these measures in explaining age-related variability in brain structure, and their relation to fractal dimensionality, using two open-access magnetic resonance imaging (MRI) datasets with a combined sample size of over 600 healthy adults across the lifespan.

## 2. Research Design and Methods

### 2.1. Datasets

**Sample 1 (OASIS)** consisted of 314 healthy adults (196 females), aged 18-94, from the publicly available Open Access Series of Imaging Studies (OASIS) cross-sectional dataset (Marcus et al., 2007; http://www.oasis-brains.org). Participants were recruited from a database of individuals who had (a) previously participated in MRI studies at Washington University, (b) were part of the Washington University Community, or (c) were from the longitudinal pool of the Washington University Alzheimer Disease Research Center. Participants were screened for neurological and psychiatric issues; the Mini-Mental State Examination (MMSE) and Clinical Dementia Rating (CDR) were administered to participants aged 60 and older. To only include healthy adults, participants with a CDR above zero were excluded; all remaining participants scored 25 or above on the MMSE. Multiple T1 volumes were acquired using a Siemens Vision 1.5 T with a MPRAGE sequence; only the first volume was used here. Scan parameters were: TR=9.7 ms; TE=4.0 ms; flip angle=10°; voxel size=1.25×1×1 mm. Volumetric and fractal dimensionality analyses from the OASIS dataset were previously reported in Madan and Kensinger (2017a).

**Sample 2 (DLBS)** consisted of 315 healthy adults (198 females), aged 20-89, from wave 1 of the Dallas Lifespan Brain Study (DLBS), made available through the International Neuroimaging Data-sharing Initiative (INDI; Mennes et al., 2013) and hosted on the the Neuroimaging Informatics Tools and Resources Clearinghouse (NITRC; Kennedy et al., 2016) (http://fcon1000.projects.nitrc.org/indi/retro/dlbs.html). Participants were screened for neurological and psychiatric issues. No participants in this dataset were excluded. All participants scored 26 or above on the MMSE. T1 volumes were acquired using a Philips Achieva 3 T with a MPRAGE sequence. Scan parameters were: TR=8.1 ms; TE=3.7 ms; flip angle=12°; voxel size=1×1×1 mm. See Kennedy et al. (2015) and Chan et al. (2014) for further details about the dataset.

### 2.2. Segmentation and volumetric analyses

All structural MRIs were processed using FreeSurfer 5.3.0 on a machine running CentOS 6.6 (Fischl, 2012; Fischl & Dale, 2000; Fischl et al., 2002). FreeSurfer’s standard pipeline was used (i.e., `recon-all`). Segmented volumes from all participants were visually inspected but no manual edits were made. Data from two additional participants were excluded from Sample 1 (OASIS) due to poor reconstructions; none were excluded from Sample 2 (DLBS). Visual inspections were conducted using Mindcontrol (Keshavan et al., in press).

FreeSurfer’s segmentation procedure produces labels for the subcortical structures within a common segmentation volume (Fischl et al., 2002, 2004). Volumes for subcortical structures were obtained directly from FreeSurfer. Validation studies have shown that this automated segmentation procedure corresponds well with manual tracing (e.g., Fischl et al., 2002; Keller et al., 2012; Lehmann et al., 2010). FreeSurfer has been used in a large number of studies investigating age-differences in subcortical structures (e.g., Inano et al., 2013; Long et al., 2012; Madan & Kensinger, 2017a; Potvin et al., 2016; Tamnes et al., 2013; Walhovd et al., 2005, 2011; Yang et al., 2016). Intracranial volume (ICV) was also estimated using FreeSurfer (Buckner et al., 2004), which has also been shown to correspond well with manual tracing (Sargolzaei et al., 2015).

### 2.3. Fractal dimensionality (FD) analyses

The complexity of each structure was calculated using the calcFD toolbox (Madan & Kensinger, 2016; http://cmadan.github.io/calcFD/). This toolbox calculates the ‘fractal dimensionality’ of a three-dimensional (3D) structure, and is specifically designed to use intermediate files from the standard FreeSurfer analysis pipeline, here `aparc.a2009s+aseg.mgz.` The toolbox has previously been used with parcellated cortical and subcortical structure, as well as validated using test-retest data (Madan & Kensinger, 2016, 2017a,b).

We use fractal dimensionality as a measure of the complexity of a 3D structure, i.e., a subcortical structure. Unlike volume, which corresponds to the ‘size’ of any 3D structure, fractal dimensionality measures shape information and is scale invariant (Madan & Kensinger, 2016, 2017a). In other words, two structures of the same shape could be different in size and still have the same fractal dimensionality. In fractal geometry, several approaches have been proposed to quantify the ‘dimensionality’ or complexity of natural structures; the approach here calculates the Minkowski-Bouligand or Hausdorff dimension (Kennedy & Lin, 1986; Mandelbrot, 1967). See Madan and Kensinger (2016, 2017a) for further details on applying fractal dimensionality to characterize cortical and subcortical structures.

### 2.4. Morphological measures of interest

A series of steps were necessary to calculate the four shape measures used here. The voxel-based segmented structure was read into MATLAB from FreeSurfer’s ‘aseg’ volume. The triangulated surface mesh (‘isosurface’) for each subcortical structure was then estimated using the marching cubes algorithm (Lorensen & Cline, 1987). The mesh was subsequently smoothed and re-parameterized relative to a sphere using an isotropic heat diffusion algorithm, as implemented by Chung (Chung 2013, 2014; Chung et al., 2008, 2010), over five iterations. A first-order ellipsoid was then fit to the surface vertices to determine a registration of the structure to standardized orientation—rather than being oriented based on native space (Cong et al., 2014; Huang et al., 2007; Shen et al., 2007, 2009), by means of a principal components analysis. The orientation of each structure was then rotated such that the long-axis corresponded to the major axis of the first-order ellipsoid. With the structure parameterized relative to a sphere, the vertex coordinates of the mesh were parameterized consistently with Euler angle conventions, with *θ* (theta) corresponding to the position relative to the poles of sphere [0, π] (akin to latitude) and *φ* (phi) corresponding to the position along the equator [0, 2π] (akin to longitude), as shown in Figure 1. This re-parameterization of a 3D closed surface to a sphere was conducted consistently with prior work (Brechbühler et al., 1995; Chung et al., 2008; Shen & Makedon, 2006; Staib & Duncan, 1996). As noted earlier, the poles of the coordinates were defined based on the long-axis of the structure, irrespective of the orientation of the structure within the brain, based on the fitted first-order ellipsoid (see Shen & Makedon, 2006).

For three of the measures (all except for sphericity), values were subsequently log-transformed. Additionally, since potential hemispheric differences were not of interest here, measures were averaged into a single value per structure and individual, collapsing across hemisphere.

#### 2.4.1. Surface-to-volume ratio (SV)

Here we simply divided the surface area of the constructed surface mesh of the structure by the volume of the structure as a coarse measure of the compactness of the structure.

#### 2.4.2. Sphericity (Sph)

The ratio of the surface area of a sphere with the same volume as the structure, relative to the actual surface area of the structure (Wadell, 1935), defined as:

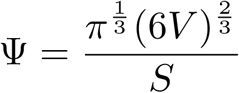

where *V* represents the volume, *S* represents the surface area, and *Ψ* (Psi) represents the structure’s sphericity.

#### 2.4.3. Long-axis curvature (LAc)

A three-dimensional (3D) smoothing spline was fit to the mean vertex coordinates of the structure, based on grouping vertices into 50 bins, with bins based on percentiles of *θ* values. The length of this spline in 3D space served as the length of the mean meridian of the structure. A second line was calculated as the straight line between the start and end points of the spline. As such, if the points along the mean-meridian spline lay perfectly along this straight line, the structure would have no long-axis curvature.

#### 2.4.4. Surface texture (Tex)

Using the Euler angle parameterization of the structure, we computed a weighted spherical harmonics (SPHARM) representation to characterize the structure across different topological frequencies (Chung 2013, 2014; Chung et al., 2008, 2010). This approach also attenuates the Gibbs phenomenon (ringing artifact) that is otherwise introduced by fitting Fourier series to discontinuous data. Here we calculated the surface texture as the ratio between the surface areas of a detailed mesh that includes high-frequency topological properties (maximum SPHARM degree 30) and a relatively smooth surface that only characterizes low-frequency topology (maximum SPHARM degree 5). A degree of 5 was selected as an appropriate threshold for low-frequency shape characteristics based on the surfaces examined in prior studies (Chung, 2013; Chung et al., 2008; Madan & Kensinger, 2017b). Examples of these two representations for the thalamus of a representative young adult are shown in Figure 1.

### 2.5. Data analyses

Age differences in the subcortical structures was first assessed using regression models examining the relationships between age and volume (or fractal dimensionality) of the structure, with the amount of variance explained (i.e., *R^2^)* and Bayesian Information Criterion *(BIC)* as the model fitness statistic. A spline regression was used as Fjell et al. (2010, 2013) demonstrated that age-related differences in structural measures are not explained well by linear and quadratic models. A smoothing spline regression was used (smoothing parameter set to 0.1), and in the case of several structural measures (i.e., the ‘Shape’ model, described below), a multiple smoothing-spline regression procedure was used, as implemented in the Prism toolbox (Madan, 2016). All regression models reported controlled for the main effect of sex. All regressions with age were conducted such that the age was the dependent variable, rather than the independent variable (i.e., unlike Madan & Kensinger, 2017a; Walhovd et al., 2011). The ‘Shape’ model is the result of a multiple spline regression including the four distinct shape measures: surface-to-volume ratio (SV), sphericity (Sph), long-axis curvature (LAc), and shape texture (Tex). A set of regression models combining measures across all four subcortical structures was also included to provide both an over-arching set of regression models across the structures, as well as show the independence vs. collinearity of the age-related differences across structures.

Volume was ICV-corrected prior to conducting the regression analyses. ICV-corrected measurements were calculated as the residual after the measure was regressed for ICV (as in Madan & Kensinger, 2017a; Walhovd et al., 2011). All shape measures— fractal dimensionality, surface-to-volume ratio, sphericity, long-axis curvature, and shape texture—are scale invariant and thus were not ICV-corrected.

For each regression model, we report both *R^2^,* with age (or fractal dimensionality) as the dependent measure, as well as the Bayesian Information Criterion (BIC). *BIC* is a model fitness index that includes a penalty based on the number of free parameters (Schwarz, 1978). Smaller *BIC* values correspond to better model fits. By convention, two models are considered equivalent if *ΔBIC* < 2 (Burnham & Anderson, 2004). As *BIC* values are based on the relevant dependent variable, *ΔBIC* values are reported relative to the best-performing model (i.e., *ΔBIC* = 0 for the best model considered).

For the models explaining age-related variability, since they all have the same dependent variable, *ΔBIC* values can be compared across all subcortical structures and measures. However, for the models with a subcortical structure‘s fractal dimensionality (FD) as the dependent measure, the *ΔBIC* values *cannot* be compared directly. Bestfitting models for each structure (thalamus, putamen, caudate, hippocampus, combined), sample (OASIS, DLBS), and dependent variable (age, FD) are shown in bold in Table 1.

**Table 1.**
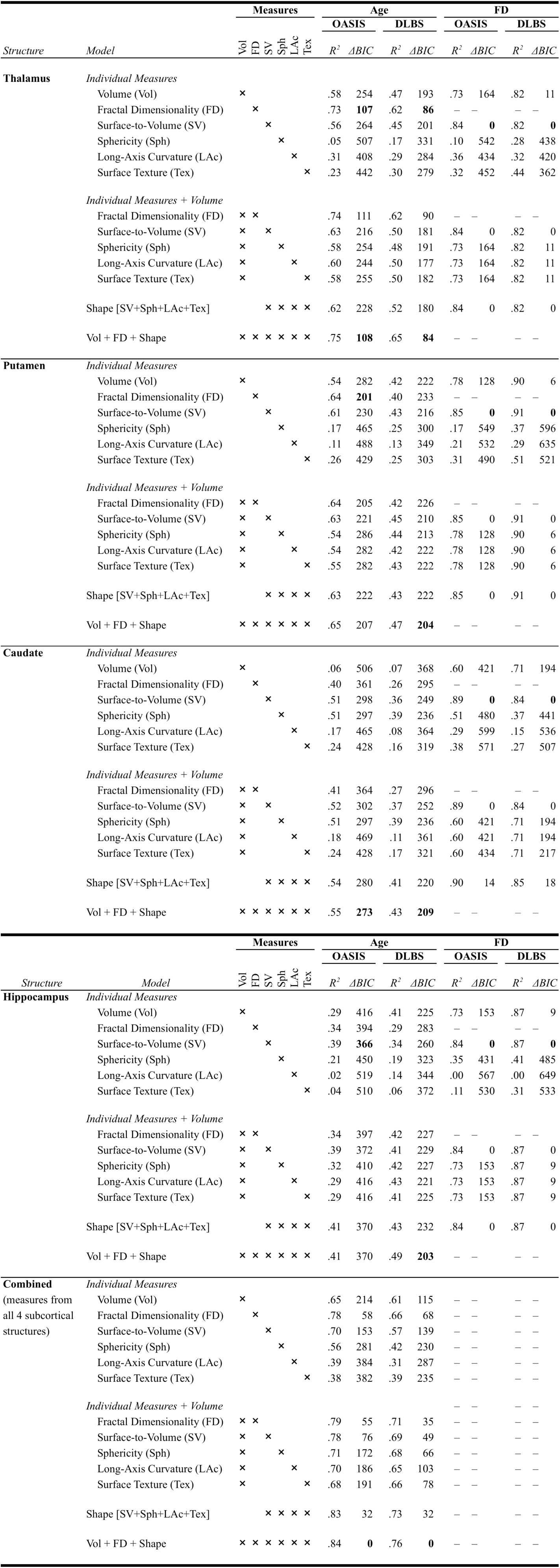
Hierarchical regression analysis of age and fractal dimensionality, for each of the structures and morphological measures considered. Best-fitting models for each structure (thalamus, putamen, caudate, hippocampus, combined), sample (OASIS, DLBS), and dependent variable (age, FD) are shown in **bold**.

Equivalent *R^2^* and *ΔBIC* values for models that include more than one measure indicate Prism algorithm (based on relevance vector regression [RVR]; Tipping, 2000) selected the same subset of measures, based on the inherent feature selection (i.e., automatic relevance determination) in RVR. E.g., for the regression models with FD as the dependent variable, if volume was a relatively good predictor, models that included volume along with a shape measure could be based only on the volume measure after the feature selection. As such, these models will all yield an identical output as the volume-only model, since the additional measure was removed. In these cases, only the simpler model is shown in bold in Table 1.

## 3. Results

Figure 2 and Table 1 show how well each of the morphological measures was able to index age-related differences in the subcortical structures. Surprisingly, the most coarse shape measure included here, surface-to-volume ratio (SV), performed the best out of the four distinct shape measures. Moreover, the aggregate ‘Shape’ model that included all four of the shape measures generally performed only slightly better than the surface-to-volume ratio alone. In both samples, the surface-to-volume ratio explained more age-related variability in brain structure than fractal dimensionality for the caudate and hippocampus. Regression models including shape measures as well as volume (see Table 1), further demonstrate that shape-related characteristics were beneficial measures of age-related differences in subcortical structure beyond volumetry.

Sphericity performed more poorly than surface-to-volume ratio in nearly all cases, despite being closely related measures. Relatedly, the long-axis curvature performed more poorly than expected, together indicating that shape information related to the elongation of the structure is not particularly useful in understanding age-related differences in subcortical structure. Higher-frequency spatial information, i.e., shape texture, also did not seem be very informative either, despite artifactual reasons that it may have been useful (e.g., head motion would lead to smoother estimates of segmented structures, older adults are known to have increased head motion; see Madan & Kensinger, 2016, for a more detailed discussion).

When the four distinct shape measures were combined with fractal dimensionality and volume (the gray bar), gains were relatively small relative to fractal dimensionality alone. However, this result is in-line with the primary goal of the study—to better characterize the structural properties that fractal dimensionality was sensitive to, using more interpretable measures of a structure’s shape. In this vein we were successful, the aggregate Shape model accounted for 80-90% of the variance in fractal dimensionality in all cases (i.e., for each subcortical structure and sample; see Table 1). The principle contributor in explaining age-related variability in fractal dimensionality was the surface-to-volume ratio measures, convergent with this measure being the most sensitive to age-related differences, of the four shape measures.

With regards to individual subcortical structures, we found that fractal dimensionality continued to be indicative of age-related differences in thalamus, even beyond the distinct shape measures considered here. Age-related differences in the two structures with the most elongation, the caudate and hippocampus, were not particularly well explained by any of the shape measures. At least, however, the shape measures did provide a significant improvement over volume, which was relatively unaffected by age. Smoothing spline fits for volume, fractal dimensionality, and surface-to-volume ratio are shown in Figure 3. These spline fits show that many middle-age adults have comparable volume and fractal dimensionality—for the caudate and hippocampus—to young adults, which is likely related to the poorer age-related differences observed here.

**Figure 3.**
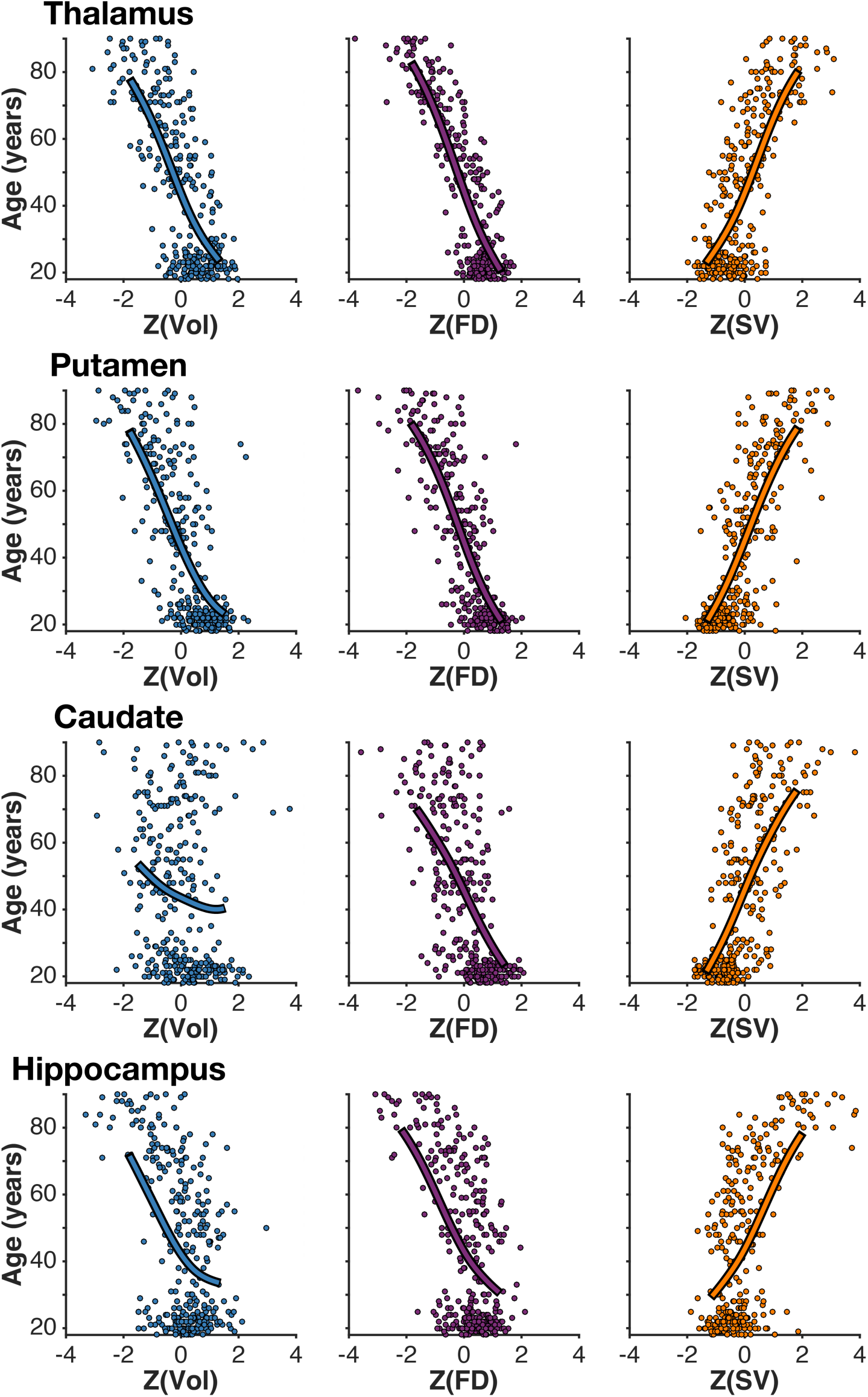
Smoothing spline age-structure fits for the OASIS sample. X-axis values represent z-scored volume (Vol), fractal dimensionality (FD), and surface-to-volume ratio (SV). Each dot represents an individual.

## 4. Discussion and Implications

Fractal dimensionality appears to be a structural measure high in reliability (Madan & Kensinger, 2017b), sensitive to age-related differences (Madan & Kensinger, 2016, 2017a), as well as useful in differentiating individuals with a variety of psychiatric and neurological disorders relative to healthy controls (de Miras et al., in press; King et al., 2010; Nenadic et al., 2014; Sandu et al., 2008; Thompson et al., 2005). However, this measure is borne out of mathematical principles, rather than quantifying a neurobiologically relevant biomarker directly. Here we compared the sensitivity of fractal dimensionality to age-related differences in healthy adults with four distinct shape-related measures that are more biologically relevant than fractal dimensionality: surface-to-volume ratio, sphericity, long-axis curvature, and surface texture. Though our results demonstrate that these other shape-related measures are able to explain most of the same variance as fractal dimensionality, we nonetheless suggest that fractal dimensionality is the more useful *single* measure, as it simultaneously accounts for these shape-related characteristics and also works as a general purpose measure of structural complexity (see Madan & Kensinger, 2016). Nonetheless, the current results indicate that surface-to-volume ratio is also a particularly useful biological marker of age-related differences in subcortical structures and should be considered in future studies of age-related structural differences. These results lay the foundation for future *ex vivo* histological research to examine how aging effects the microstructure of subcortical structures.

Here we demonstrate that shape-related measures can be used as robust biological markers of aging using a computational neuroanatomy framework. While fractal dimensionality performed well, the four distinct measures of shape-related characteristics were also sensitive to age, particularly surface-to-volume ratio. Furthermore, the current approach is in-line with the emerging literature on ‘radiomics’ (Adduru et al., in press; Gillies et al., 2016; Lambin et al., 2012, in press; Parekh & Jacobs, 2016; Yip & Aerts, 2016), the use of high-throughput automatic quantitative imaging analyses to calculate structural features related to the shape of brain structures from radiological images, as well as further demonstrates the benefits of open-access data for brain morphology research (see Madan, 2017, for an in-depth discussion). The current findings clarify the age-related differences in the shape, not just volume, of subcortical structures in the brain and provide strong evidence for additional biological markers of aging.

## Acknowledgements

I would like to thank Elizabeth Kensinger for feedback on an earlier version of the manuscript. This work was supported by a fellowship from the Canadian Institutes of Health Research [FRN-146793].

MRI data used in the preparation of this article were obtained from several sources, data were provided in part by: (1) the Open Access Series of Imaging Studies (OASIS; Marcus et al., 2007); and (2) wave 1 of the Dallas Lifespan Brain Study (DLBS) led by Dr. Denise Park, and distributed through INDI (Mennes et al., 2013) and NITRC (Kennedy et al., 2016).

